# The fatty liver disease-causing protein PNPLA3-I148M alters lipid droplet-Golgi dynamics

**DOI:** 10.1101/2023.10.13.562302

**Authors:** David J. Sherman, Lei Liu, Jennifer L. Mamrosh, Jiansong Xie, John Ferbas, Brett Lomenick, Mark S. Ladinsky, Rati Verma, Ingrid C. Rulifson, Raymond J. Deshaies

## Abstract

Non-alcoholic fatty liver disease (NAFLD), recently renamed metabolic dysfunction-associated steatotic liver disease (MASLD), is a progressive metabolic disorder that begins with aberrant triglyceride accumulation in the liver and can lead to cirrhosis and cancer. A common variant in the gene *PNPLA3*, encoding the protein PNPLA3-I148M, is the strongest known genetic risk factor for MASLD to date. Despite its discovery twenty years ago, the function of PNPLA3, and now the role of PNPLA3-I148M, remain unclear. In this study, we sought to dissect the biogenesis of PNPLA3 and PNPLA3-I148M and characterize changes induced by endogenous expression of the disease-causing variant. Contrary to bioinformatic predictions and prior studies with overexpressed proteins, we demonstrate here that PNPLA3 and PNPLA3-I148M are not endoplasmic reticulum-resident transmembrane proteins. To identify their intracellular associations, we generated a paired set of isogenic human hepatoma cells expressing PNPLA3 and PNPLA3-I148M at endogenous levels. Both proteins were enriched in lipid droplet, Golgi, and endosomal fractions. Purified PNPLA3 and PNPLA3-I148M proteins associated with phosphoinositides commonly found in these compartments. Despite a similar fractionation pattern as the wild-type variant, PNPLA3-I148M induced morphological changes in the Golgi apparatus, including increased lipid droplet-Golgi contact sites, which were also observed in I148M-expressing primary human patient hepatocytes. In addition to lipid droplet accumulation, PNPLA3-I148M expression caused significant proteomic and transcriptomic changes that resembled all stages of liver disease. Cumulatively, we validate an endogenous human cellular system for investigating PNPLA3-I148M biology and identify the Golgi apparatus as a central hub of PNPLA3-I148M-driven cellular change.

**Significance Statement:** Fatty liver disease affects nearly a quarter of the world’s population and has both environmental and genetic risk factors. A mutation in the gene *PNPLA3* that converts Ile 148 to Met is the strongest known genetic risk factor for developing fatty liver disease. Using a series of techniques to track endogenous PNPLA3 and PNPLA3-I148M biogenesis and localization, we reveal new insights into how the mutation changes cellular dynamics. Although previous reports focus on its role on lipid droplets, we reveal that PNPLA3-I148M also functions at the Golgi apparatus, an organelle critical for protein transport into and out of the cell and lipid signaling. PNPLA3-I148M causes altered Golgi morphology and drives changes reminiscent of liver disease.

## Introduction

Metabolic dysfunction-associated steatotic liver disease (MASLD)^†^ is a burgeoning public health threat, estimated to affect approximately one quarter of the world’s population and growing in prevalence similarly to obesity and type 2 diabetes (1, 2). MASLD represents a spectrum of histological changes, starting with fat accumulation in the liver (i.e., steatosis) and, in some individuals, progresses to an inflammatory condition known as metabolic dysfunction-associated steatohepatitis (MASH), followed by cirrhosis, and possibly hepatocellular carcinoma. Despite the global threat it poses, the biological underpinnings of the disease are unclear. A major advance in understanding the disease came with the identification of a nonsynonymous variant (rs738409[G]) in the gene *PNPLA3* (patatin-like phospholipase domain-containing protein 3) which is strongly associated with increased hepatic fat (3). This mutation in *PNPLA3* encodes an isoleucine-to-methionine substitution at position 148 (PNPLA3-I148M). The presence of this variant allele increases both the risk and severity across the entire disease spectrum, making it the strongest known genetic determinant of MASLD to date (4–8).

PNPLA3 is a member of a diverse protein family found in organisms ranging from bacteria to humans (9). Receiving its name from patatin, a nonspecific lipid acyl hydrolase found in potatoes, the patatin-like phospholipase (PNPLA) domain contains an α/β/α sandwich fold and a conserved catalytic dyad (Ser-Asp) lipase motif. PNPLA family members vary in their cellular localizations and have biological functions that include host colonization, membrane maintenance, and triglyceride metabolism (9, 10). Despite the significance of PNPLA3-I148M to human health, details about the function of wild-type PNPLA3 and the mechanism(s) by which PNPLA3-I148M causes pathogenesis remain elusive. Whereas deletion or overexpression of the wild-type variant in mice does not cause steatosis, transgenic PNPLA3-I148M overexpression, or knock-in of the mutation in mice challenged with a high-sucrose diet, cause steatosis (11–14). In human induced pluripotent stem cells and organoid systems, PNPLA3-I148M elicits an intermediate phenotype between the wild-type and complete *PNPLA3* knockout (15, 16). Reducing PNPLA3-I148M protein levels in the liver, including with I148M-specific siRNA, reduces fatty liver disease phenotypes, supporting a specific role for PNPLA3-I148M in driving MASLD (17–19). Purified wild-type PNPLA3 has been shown to possess triglyceride hydrolase or lysophosphatidic acid acyltransferase (LPAAT) activity, with the mutant variant losing the hydrolase activity or showing elevated LPAAT activity (20–25). However, *in vivo* and cellular data suggest that the conserved enzyme active site is not required for I148M-driven disease. In sum, PNPLA3-I148M can best be described as a neomorph, with the gene product possessing a novel molecular function. A model that has been proposed is that PNPLA3-I148M sequesters CGI-58/ABHD5, a cofactor for the triglyceride hydrolase ATGL/PNPLA2, impairing triglyceride hydrolysis on cytosolic lipid droplets (LDs) and thereby driving steatosis (26). More studies are needed to demonstrate that this effect is specific to the mutant variant and how this function accounts for the pleiotropic effects caused by PNPLA3-I148M. PNPLA3 was originally described as a transmembrane protein that can fractionate with cytosolic LDs and intracellular membranes (24, 25, 27, 28). Cellular studies with overexpressed PNPLA3 and PNPLA3-I148M suggested that the proteins either localize to the endoplasmic reticulum (ER) or are generally cytosolic in nature (26, 28). Considering the significant hydrophobicity of the proteins (containing nearly 50% amino acids with hydrophobic uncharged side chains), it is likely that PNPLA3 and PNPLA3-I148M would need to partition into a membrane structure if not bound to a chaperone or LD (*vide infra*). Residues 42-62 have been manually annotated as a type II signal anchor sequence (uniprot.org), with the majority of the PNPLA3 polypeptide predicted to be in the ER lumen (**Fig. S1A**). The ER is a critical compartment for lipid metabolism and LD biogenesis, with known pathways for ER-to-LD protein trafficking (29, 30), making it a plausible localization site for PNPLA3. However, a model in which PNPLA3 is transmembrane (either with a signal anchor sequence or multiple transmembrane helices) is difficult to rationalize because it would require major structural rearrangements to associate with LDs, which are unique organelles in that they contain a core of neutral lipids surrounded by a phospholipid monolayer (29, 31). Therefore, LDs lack a lumenal compartment for the soluble extracytosolic portions of transmembrane proteins. Nevertheless, a transmembrane orientation for PNPLA3 has not been formally disproven, nor have the localizations of the endogenous human wild-type and I148M variants been determined. Disentangling the site(s) of activity of PNPLA3-I148M is important for understanding its function in lipid homeostasis.

Several studies to investigate PNPLA3-I148M function to date have been conducted in mice, yet the mouse and human orthologs differ in their tissue distribution and in their primary sequences and lengths, confounding extrapolations to human protein mechanism and function (8). Given that MASLD is a major emerging public health crisis and PNPLA3-I148M is the single most important human genetic determinant of the disease, we have set out to understand in detail the biochemistry of the wild-type and mutant protein variants and their impact on cellular biology. In this work, we sought to dissect the biogenesis of the human PNPLA3 and PNPLA3-I148M proteins. Using paired human hepatoma cell lines expressing PNPLA3 and PNPLA3-I148M at endogenous levels, we identify the Golgi apparatus as a central node in PNPLA3-I148M-driven cellular change and characterize alterations in LD-Golgi dynamics that translate to primary human hepatocytes.

## Results

### Constitutive endogenous expression of PNPLA3-I148M increases cellular LD content

To investigate the biogenesis and cellular effects of endogenously expressed human PNPLA3 and PNPLA3-I148M, we generated a paired set of Hep3B cells using CRISPR. Hep3B is a hepatoma cell line that expresses endogenous wild-type PNPLA3, whereas several typical human hepatoma cell lines express endogenous PNPLA3-I148M. We gene-edited the native *PNPLA3* locus in Hep3B cells to make a homozygous PNPLA3-I148M (I148M) cell line (**Fig. 1A**). Because PNPLA3 is a low-abundance protein and we have been unable to track the endogenous gene product with existing antibodies, we introduced a HiBiT tag at the C-terminus of PNPLA3 in both WT and I148M cells (32, 33). We confirmed homozygous tagging in the WT and I148M cells. In Hep3B knock-in cells, mRNA levels of PNPLA3 and PNPLA3-I148M were equal, yet the steady-state level of PNPLA3-I148M was ∼ 3-4 times greater than that of the WT protein (**Fig. 1B**). This recapitulates what has been reported in mouse models (14).

**Figure 1.**
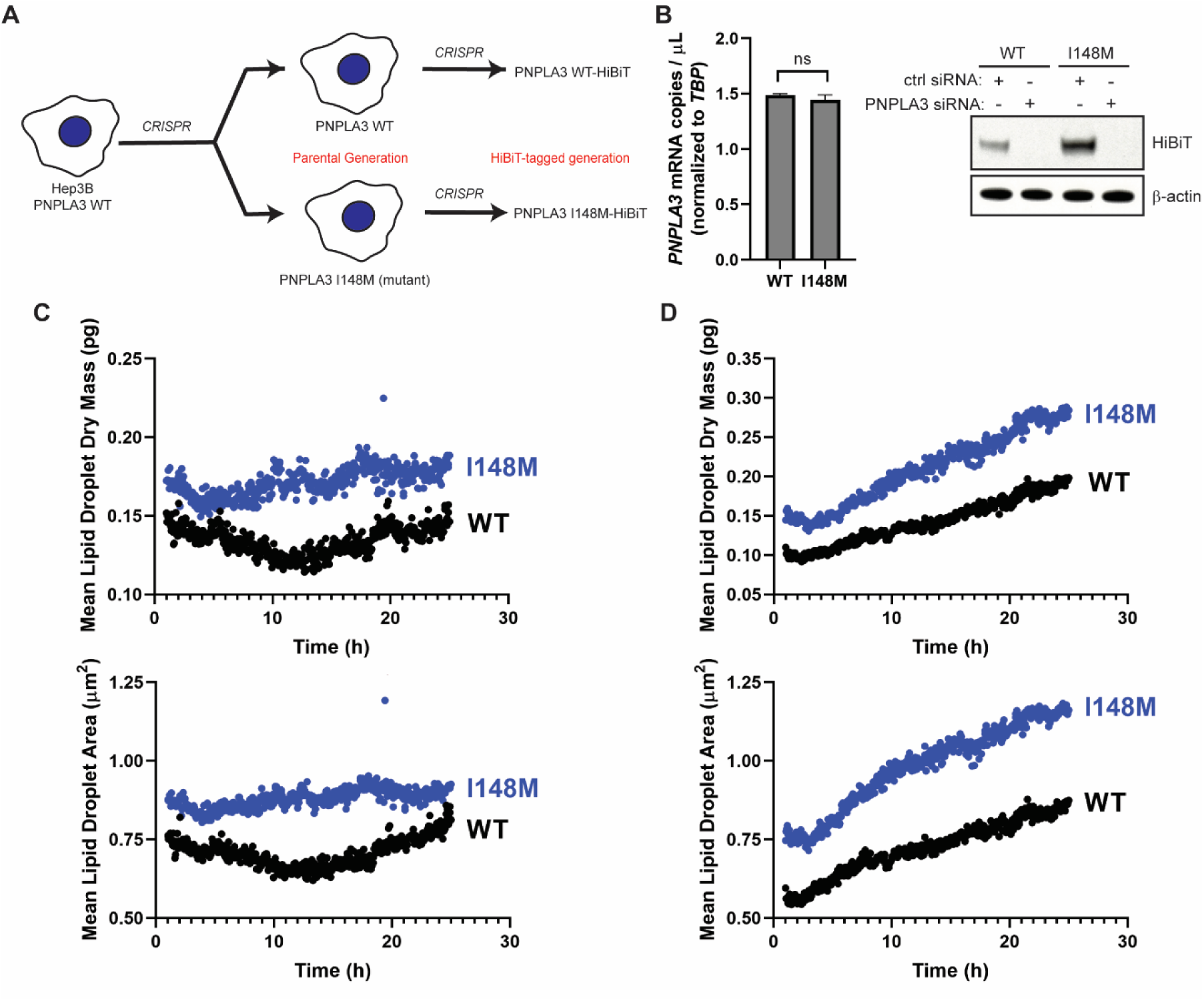
Constitutive endogenous expression of PNPLA3-I148M increases the LD content of hepatoma cells. (A) *PNPLA3-I148M* was introduced into Hep3B cells using CRISPR. Parental Hep3B cells (WT and I148M) were further modified to introduce a C-terminal HiBiT tag at the endogenous locus. (B) *PNPLA3* expression in the parental cells (left) was analyzed by digital PCR (*p =* 0.0972, determined by Student’s *t*-test, n = 3). Protein expression (right) was analyzed by HiBiT detection in the tagged cell lines treated with a negative control siRNA or PNPLA3-specific siRNA. (C,D) Mean lipid droplet content (upper) and mean lipid droplet area (lower) were quantified from live-cell label-free imaging using the Nanolive 3D Cell Explorer microscope. Cells were treated with vehicle (C) or with 200 µM oleic acid (D). Images were taken every 3 min for 24 h, starting 1 h after addition of oleic acid. Quantification was done using Eve software.

To evaluate the impact of PNPLA3-I148M expression on the LD content of hepatoma cells, we performed live-cell, label-free imaging using Nanolive, a 3D holotomographic microscopic platform that allows for LD identification by their high refractive index values (34, 35). Even under basal conditions, I148M cells had ∼ 1.4 times greater mean LD dry mass and area than WT cells (**Fig. 1C**). When cells were treated with oleic acid, there was a steady increase in LD content throughout a 24-hour time course, with at least ∼ 1.6 times greater mean LD dry mass and area in I148M cells than in WT cells (**Fig. 1D**). Mean LD dry mass and area increased in I148M cells at ∼ 1.3 times the rate of WT cells. This mirrors reports that expression of PNPLA3-I148M is enough to drive increases in LD content in the mouse liver and in overexpressing cell lines (13, 14, 26, 36).

### PNPLA3-I148M fractionates with the Golgi and alters Golgi structure

PNPLA3 contains several stretches of hydrophobic amino acids that are of the right length to span a membrane, three of which are in the PNPLA domain (**Fig. S1A**). If PNPLA3 is in fact an integral membrane protein that localizes to the ER, then it must contain an ER-targeting signal sequence. To investigate ER membrane targeting and association, we employed a biochemical system to investigate the process *in vitro*. Human PNPLA3 and PNPLA3-I148M with C-terminal hemagglutinin (HA) epitope tags were translated in rabbit reticulocyte lysate in the presence of ^35^S-methionine. In some reactions, canine rough microsomal membranes (cRMs) were included (co-translationally) or were added after termination of translation (post-translationally). cRMs were isolated by centrifugation, washed with sodium carbonate to remove peripherally associated proteins, and re-isolated. The majorities of PNPLA3-HA and PNPLA3-I148M-HA were found in the supernatant fraction (**Fig. S1B**), with ∼ 10% of the wild-type protein and < 5% of the I148M variant found stably inserted in the cRMs post-translationally (**Fig. S1C**). In contrast, a construct with a deletion of the hydrophobic stretch of amino acids 42-62 was found predominantly in the original supernatant fraction, not bound to cRMs. Due to the small fraction of PNPLA3-HA stably associated with cRMs, the lack of association of the Δ42-62 construct does not provide conclusive evidence that amino acids 42-62 constitute a signal anchor sequence. Membrane-associated PNPLA3-HA and PNPLA3-I148M-HA were completely digested by Proteinase K (**Fig. S1D**), inconsistent with the prediction that PNPLA3 contains a significant protease-protected, ER-lumenal C-terminal domain (**Fig. S1A**). This lack of protease protection was also true for microsomes isolated from HEK293T cells expressing PNPLA3 (**Fig. S1E**). Although the protease protection experiments suggest that PNPLA3-HA does not contain an ER-lumenal portion, we tested the hypothesis that parts of PNPLA3 could be inside the ER by treating detergent-solubilized cRMs with Endoglycosidase H (Endo H), which cleaves asparagine-linked mannose-rich oligosaccharides that would be added to PNPLA3 if its consensus N-linked glycosylation sites were inside the microsomal lumen. PNPLA3-HA and PNPLA3-I148M-HA did not show gel migration shifts by SDS-PAGE when digested with Endo H, nor were the N- and C-termini glycosylated when an opsin tag containing a consensus glycosylation site was added (**Fig. S1F**).

If PNPLA3 contains an ER-specific signal sequence (even if not a transmembrane signal anchor), there is conserved ER-specific protein machinery that would be required to recognize and insert it into the membrane (37). To determine if protein insertases at the ER are required for stable association of PNPLA3-HA or PNPLA3-I148M-HA in this biochemical system, we post-translationally added cRMs that were trypsin-treated (“T-cRMs”) or “mock” treated (“M-cRMs”) to *in vitro* translation reactions (38, 39). We hypothesized that if membrane insertion relied on traditional pathways at the ER, digesting the exposed membrane proteins with trypsin would eliminate protein insertion. We found that PNPLA3-HA and PNPLA3-I148M-HA could still be inserted into T-cRMs (**Fig. S1G**). Taken together, these experiments suggest that PNPLA3 and, to a lesser extent, PNPLA3-I148M, can insert into ER-derived membranes without membrane-resident protein insertion machinery, and the portion of each protein stably inserted into the membrane is small and does not traverse the phospholipid bilayer. This is inconsistent with traditional ER transmembrane proteins but consistent with traditional LD proteins, which often form membrane hairpins that allow for stable insertion into the outer phospholipid monolayer (40).

Next, we sought to determine where PNPLA3 and PNPLA3-I148M localize in the endogenous Hep3B cellular system. We fractionated the post-nuclear supernatant of basal-state Hep3B cells into cytosolic and crude membrane fractions. PNPLA3-HiBiT and PNPLA3-I148M-HiBiT proteins isolated entirely with crude membranes (**Fig. 2A**). This differed from perilipin-2 (PLIN2), the majority of which remained in the cytosolic fraction, consistent with its designation as a type II LD protein that is targeted to LDs from the cytosol (31). We next determined the specific membrane fraction(s) with which PNPLA3 and PNPLA3-I148M isolate. ∼ 78% of PNPLA3-HiBiT and ∼ 69% of PNPLA3-I148M-HiBiT were isolated with the Golgi membranes, with the remaining protein pools found in the “other membranes” fraction, which included the ER and plasma membrane (**Fig. 2B**).

**Figure 2.**
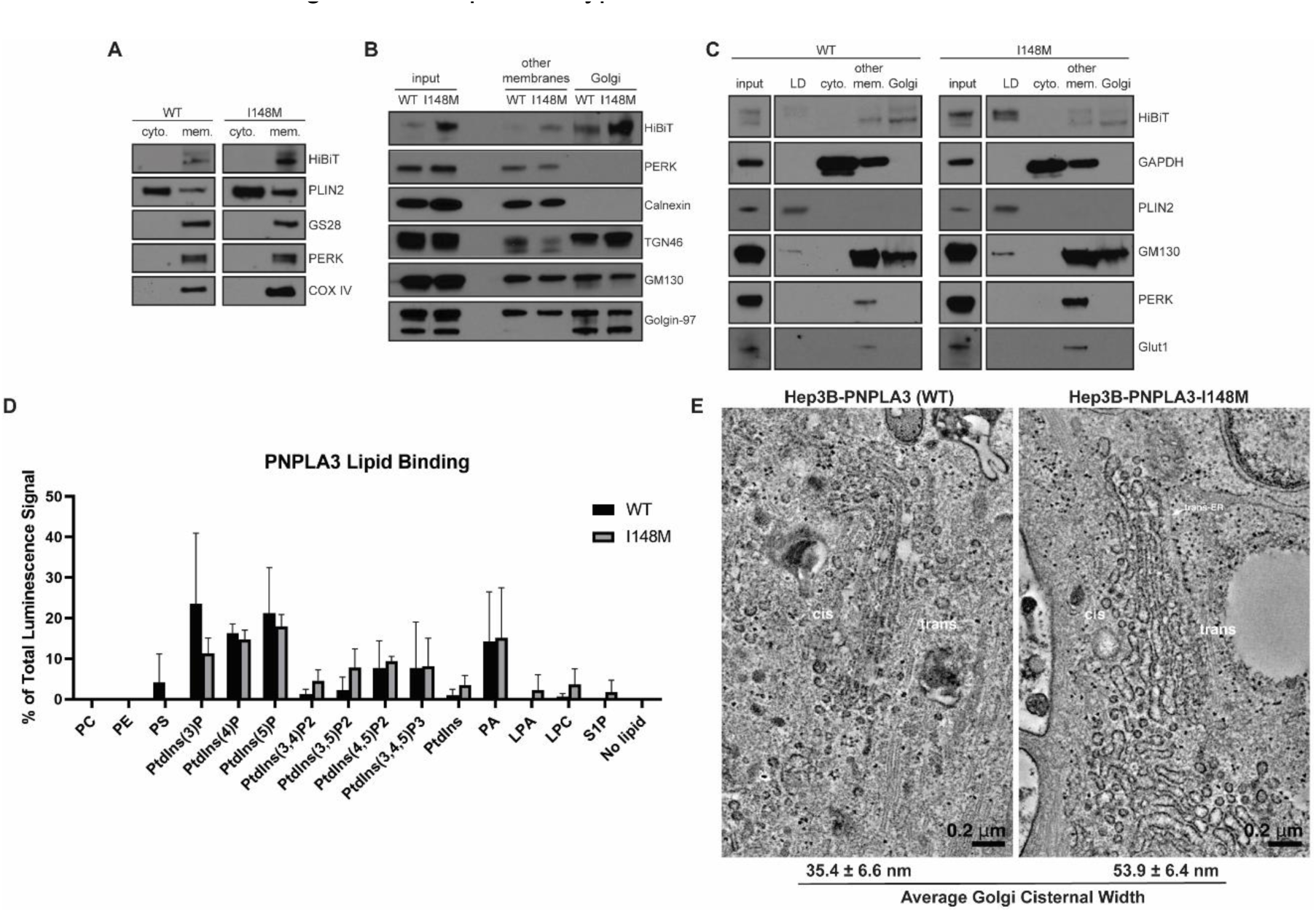
PNPLA3-I148M induces structural changes at the Golgi apparatus. (A) Basal-state Hep3B cells expressing endogenous PNPLA3 or PNPLA3-I148M were fractionated into cytosolic (“cyto.”) or total membrane (“mem.”) fractions. (B) Basal-state Hep3B cells were further fractionated to separate membrane species. “Other membranes” include non-Golgi membranes. (C) Hep3B cells were treated with 100 µM oleic acid for 16 h prior to fractionation into LD, cytosolic, “other membranes,” and Golgi membrane fractions. (D) Purified PNPLA3-His and PNPLA3-I148M-His were incubated with phosphoinositide strips containing spots with 100 pmol of different lipid species. Following washing and staining with anti-His-HRP, strips were imaged using chemiluminescence. Bar graph represents the average of three independent experiments. *p*-values indicated in Table S5. (E) Hep3B cells (WT or I148M) were treated with 100 µM oleic acid for 16 h, then fixed and imaged by transmission electron microscopy. Average cisternal widths were calculated from 20 Golgi cisternae per cell type.

When Hep3B cells were treated with oleic acid overnight to stimulate LD formation, ∼ 15% of the WT protein and ∼ 72% of I148M redistributed to the buoyant LD fractions (**Fig. 2C**), whereas ∼ 61% and ∼ 18% of the WT and I148M proteins, respectively, were found in the enriched Golgi fractions. The remaining protein pools were isolated with “other membranes.” In a separate experiment, PNPLA3-HiBiT and PNPLA3-I148M-HiBiT were found in endosomes isolated from basal-state Hep3B cells (**Fig. S2A**). While we cannot determine if the endosomal pool is derived from the “other membrane” or Golgi fractions, and how this changes with oleic acid stimulation, it is notable that other LD proteins, including Rab GTPases, can also localize to endosomes (41).

To determine how PNPLA3 and PNPLA3-I148M might associate with the secretory/endosomal pathway, we tested whether the purified proteins bind to membrane and signaling lipids commonly found in the Golgi, endosome, and plasma membrane. PNPLA3 and PNPLA3-I148M with C-terminal hexa-histidine (His) tags bound to lipid arrays spotted with different phosphoinositide species, especially PtdIns(3)P, PtdIns(4)P, and PtdIns(5)P (**Fig. 2D** **and S2B**). These phosphoinositides are enriched in the Golgi/*trans*-Golgi network and in the endosomal system (42). PNPLA3-I148M demonstrated even greater selectivity for phosphoinositide species from membrane phospholipids than the WT protein (**Table S5**). Otherwise, there were no significant differences in the lipids bound by the WT and mutant variants. In addition to phosphoinositide binding, both proteins interacted with phosphatidic acid (**Fig. 2D**). Similar results were observed using proteins purified from whole cell lysate or from the membrane fraction, with either C-terminal His- or FLAG-tags. To confirm the quality of the lipid-binding arrays, we verified that purified GST-tagged PLC-δ1 PH domain protein correctly interacted with PtdIns(4,5)P2 (**Fig. S2B**) (43, 44).

Because PNPLA3-I148M can bind phospholipids important for vesicular trafficking, signaling and membrane dynamics, we investigated if there were morphological changes in I148M cells. Hep3B cells (WT and I148M) were treated with oleic acid, fixed, and imaged using transmission electron microscopy (TEM). Whereas the ER was similar in both cell lines, the I148M cells demonstrated marked enlargement of Golgi cisternae (**Fig. 2E**). These enlarged structures had lucent interiors and could be observed among the *cis*-, *medial*-, and *trans*-Golgi network.

### Endogenous PNPLA3-I148M alters the proteomic and transcriptomic landscape of Hep3B cells

As the nexus of the secretory and endolysosomal pathways, changes in Golgi architecture can affect fundamental cellular processes. We were interested in whether constitutive expression of endogenous PNPLA3-I148M resulted in proteomic changes in the cell. Therefore, we conducted quantitative mass spectrometry on cell lysates of WT- and I148M-expressing Hep3B-HiBiT cells treated with or without oleic acid (**Fig. 3A** **and S3A**). We observed that I148M-expressing cells had both significantly increased and decreased levels of multiple proteins compared to WT cells. The effect of the PNPLA3 variant on protein abundance levels was similar between untreated cells and cells treated with oleic acid for 16 h, suggesting that the I148M variant has a distinct effect on Hep3B cells compared to oleic acid treatment (**Fig. S3B**). To ensure that these changes were not due to clonal effects or the HiBiT tag, we confirmed top increased proteins in parental cells and other HiBiT cell clones (**Fig. 3B**).

**Figure 3.**
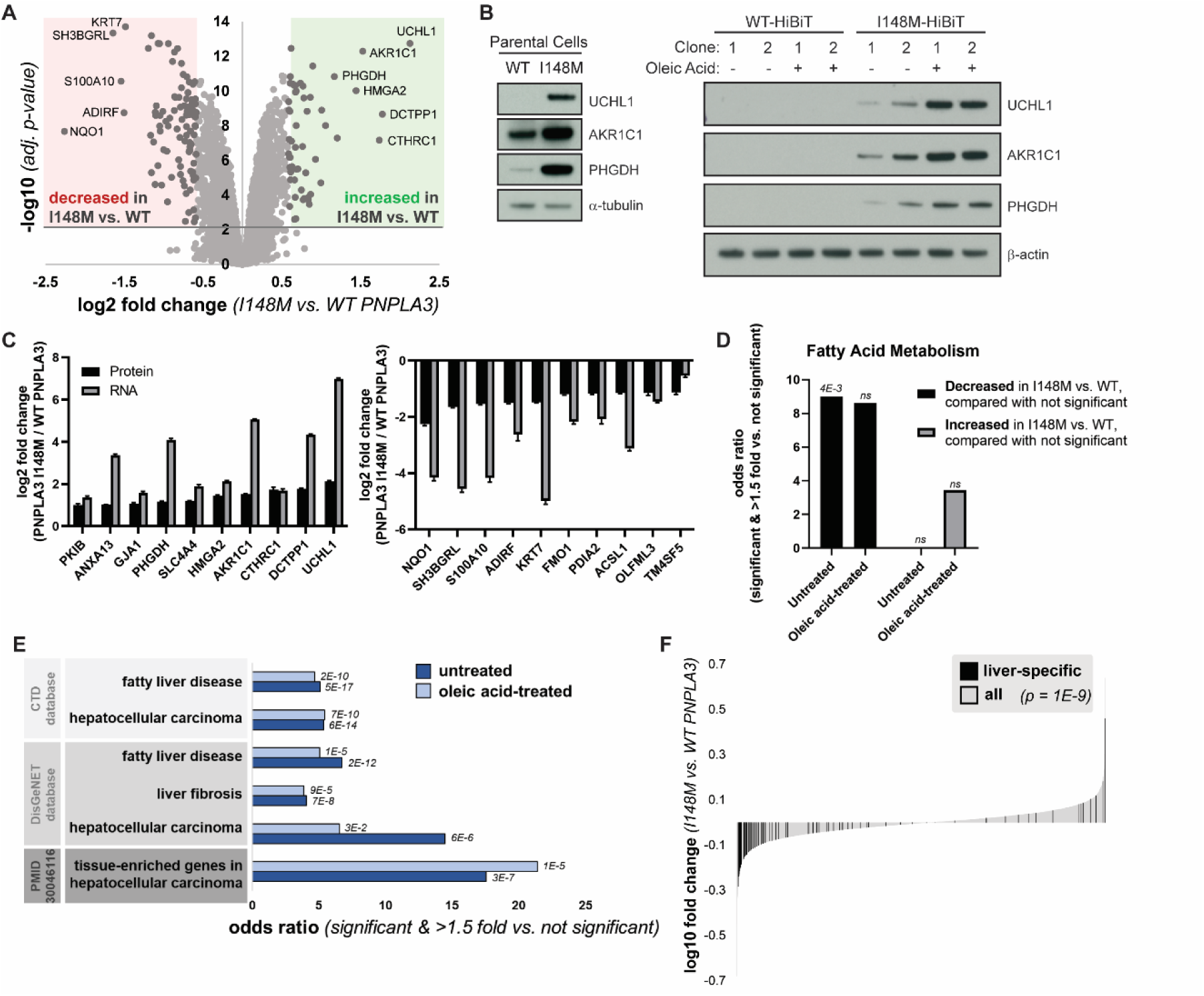
PNPLA3-I148M causes proteomic changes consistent with liver disease. (A) Volcano plot of proteins quantified by mass spectrometry in untreated cells expressing WT or PNPLA3-I148M. Red and green compartments contain proteins significantly (adj. *p*-value < 0.01) decreased or increased, respectively, by >1.5 fold in the I148M vs. WT cell lines. (B) Confirmation of top proteomic hit proteins in parental Hep3B cells and additional HiBiT clones, with and without incubation with 200 µM oleic acid overnight. (C) Log2 fold changes of top ten significantly upregulated (left) and significantly downregulated (right) proteins in untreated mass spectrometry, compared with gene expression from RNA-seq. (D) Enrichment of the biological process keyword in proteins significantly (adj. *p*-value < 0.01) decreased or increased by >1.5 fold in the I148M vs. WT PNPLA3 cell lines versus those not significantly changing. Fatty acid metabolism is the only term that is significant after multiple testing correction in at least one comparison. *p*-values indicated above the bars. (E) Proteins significantly (adj. *p*-value < 0.01) decreased or increased by >1.5 fold in the I148M vs. WT PNPLA3 cell lines were compared with those not significantly changed. Enrichment of liver disease-related terms was determined; adjusted *p*-value by Fisher’s exact test is shown. (F) Plot showing the log10 fold change for proteins quantified in the I148M vs. WT PNPLA3 untreated cell lines. Those reported (TiGER database) to be liver-specific genes are shown in black. Kolmogorov-Smirnov *p*-value testing for a difference in fold change distribution between the two protein sets is reported.

In addition to whole-cell proteomics, we performed RNA-seq on these cell lines. When comparing the proteomics and RNA-seq datasets from untreated cells, the top changing proteins in the proteomics datasets were also changing in the corresponding direction at the mRNA level (**Fig. 3C**). Therefore, these proteomic changes were due to transcriptional effects. We considered whether there were differences in biological function between proteins significantly changed in the I148M versus the WT cells, as compared with proteins not significantly changed. Proteins decreased in untreated I148M cells were more likely to be involved in fatty acid metabolism, consistent with a putative function of PNPLA3 in lipid homeostasis (**Fig. 3D**).

We also observed that proteins significantly changed in I148M versus WT cells were more likely to overlap with genes associated with fatty liver disease, liver fibrosis, and hepatocellular carcinoma (HCC) (**Fig. 3E, S3C**). Furthermore, liver-specific genes were enriched in proteins significantly changed in these cell lines (**Fig. 3F**). These genes, the majority of which were downregulated in I148M, reflect a “dedifferentiation” of hepatocytes observed in fatty liver disease and HCC (45, 46). Indeed, proteins decreased in the I148M-expressing cells were more likely to be liver-specific genes that are also downregulated in HCC (**Fig. S3D**). Therefore PNPLA3-I148M, even in the absence of exogenous lipid stimulation, is sufficient to promote an effect resembling dedifferentiation of hepatocytes associated with the development of HCC, even when evaluated in hepatoma cell lines.

### Endogenous PNPLA3-I148M increases LD-Golgi contact sites in primary human hepatocytes

Hep3B cells expressing PNPLA3-I148M have greater LD content (**Fig. 1C,D**), and qualitatively, there appeared to be more LDs in the vicinity of the Golgi compared to WT cells, as judged by TEM imaging (**Fig. S4**). To test whether there were changes in LD-Golgi contacts, Hep3B cells were fixed, permeabilized, and stained with anti-TGOLN2 (Golgi; red stain) and LipidTOX Green (LD; green stain) for visualization by confocal microscopy (**Fig. 4A**). After separately quantifying the LD and TGOLN2 staining areas using a machine learning approach, LD-TGOLN2 interactions were quantified by counting juxtaposed staining sites. Staining sites within 30 nm of each other were considered juxtaposed based on an acceptable cutoff for determining membrane contact sites (47). Consistent with the TEM imaging, mutant Hep3B cells had a greater percentage of LD stain adjacent to TGOLN2, and vice versa (**Fig. 4B**). Additionally, we qualitatively observed a more dispersed TGOLN2 signal in I148M cells.

**Figure 4.**
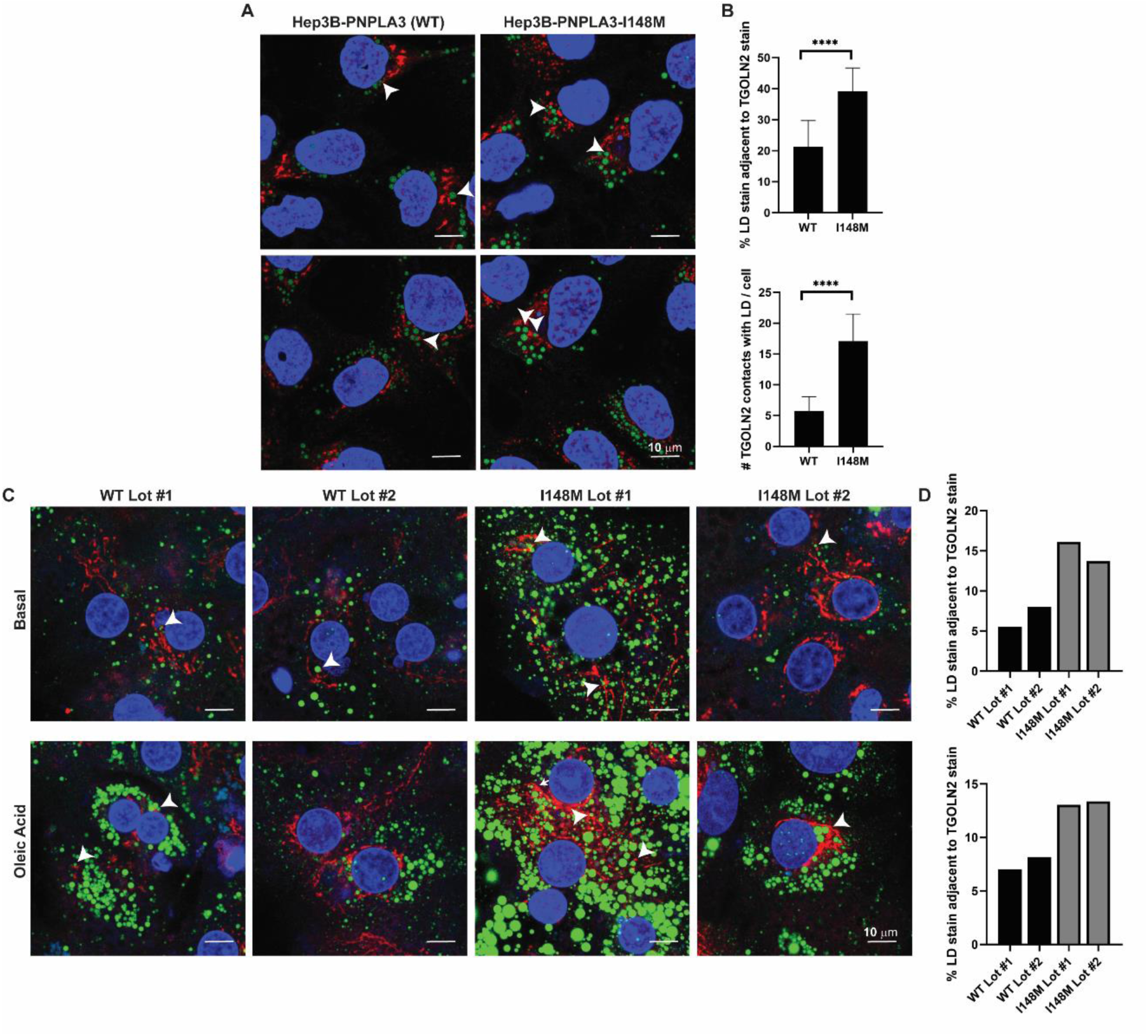
Hepatoma cells and primary hepatocytes expressing PNPLA3-I148M have more LD-Golgi contacts. (A) Hep3B cells expressing endogenous PNPLA3 or PNPLA3-I148M were fixed and stained with LipidTOX Green to label LDs (green) and TGOLN2 (red) to label the *trans*-Golgi network. White arrowheads indicate examples of LD-TGOLN2 interactions. (B) Lipid droplet-TGOLN2 contact sites in ∼ 45 cells per cell line were quantified using Imaris software. *p-*values were calculated from Student’s *t-*tests. (C) Individual lots of primary human hepatocytes expressing wild-type PNPLA3 or PNPLA3-I148M were plated, treated with vehicle or 200 µM oleic acid for 24 h, fixed and stained with LipidTOX Green to label LDs (green) and anti-TGOLN2 (red) to label the *trans*-Golgi network. White arrowheads indicate examples of LD-TGOLN2 interactions. (E) 20-40 cells of each lot and condition were quantified and LD-TGOLN2 proximity was calculated using Imaris software. Percentage of LD stain juxtaposed to TGOLN2 stain for untreated cells (top) and oleic acid-treated cells (bottom) per primary hepatocyte lot. WT (black) and I148M (gray) lots are indicated. Each bar represents the average of 11 fields of view for each primary hepatocyte lot per condition. **** *p* < 0.0001.

Considering the disease-like changes induced by endogenous PNPLA3-I148M in the hepatoma cells, we asked if endogenous expression of PNPLA3-I148M can induce similar changes in LD-Golgi dynamics in primary human hepatocytes. Primary hepatocyte lots (each lot derived from a different patient) expressing either WT PNPLA3 or homozygous for rs738409[G] were identified (**Fig. S5**). The hepatocytes were either maintained in media lacking fatty acid supplementation or treated with 200 µM oleic acid for 24 h. Then, the cells were fixed, permeabilized, and stained with anti-TGOLN2 and LipidTOX Green (**Fig. 4C**). After quantifying the LD and TGOLN2 stains separately, as in Hep3B cells, stain contact sites were quantified. Despite significant differences in LD and Golgi content between lots (even between the two PNPLA3-I148M-expressing lots), there were overall more LD-TGOLN2 contact sites in the mutant hepatocytes compared to the WT hepatocytes (**Fig. 4D**). This suggests that rs738409[G] is driving the LD-TGOLN2 contact trends despite the genetic variability in these four lots.

## Discussion

In this study, we investigated the biogenesis of PNPLA3 and PNPLA3-I148M and the cellular consequences of constitutive endogenous I148M expression in hepatoma cells. We generated a paired set of Hep3B cells expressing PNPLA3-HiBiT and PNPLA3-I148M-HiBiT and demonstrated that relative PNPLA3 abundance and cellular LD content recapitulate prior reports, including studies with overexpression cell lines and knock-in mice (**Fig. 1**) (14). Then, we sought to resolve conflicting reports on the nature of PNPLA3 localization and topology. Using an *in vitro* translation system, we could not establish that PNPLA3 is a transmembrane protein nor that it is targeted to the ER with a signal sequence (**Fig. S1**). Using a highly sensitive luminescent reporter tag, we demonstrated for the first time that endogenous PNPLA3 and PNPLA3-I148M isolate with cellular LD, Golgi, and endosomal fractions (**Fig. 2**). As a basis for such localization, we interrogated lipid binding properties of the purified WT and I148M proteins and found that they interact with phosphoinositides enriched in Golgi and endosomal membranes. Through imaging studies, we identified distinct morphological changes in the Golgi apparatus in I148M-expressing cells **(Fig. 2, S4**). To complement the morphological changes, whole-cell proteomics and RNA-seq revealed altered proteomic and transcriptomic profiles in I148M-expressing cells that are consistent with those found in the full spectrum of liver disease, supporting a role for I148M in driving all stages of MASLD (**Fig. 3**). Finally, more LD-Golgi contacts were observed in Hep3B cells and primary human hepatocytes expressing endogenous PNPLA3-I148M (**Fig. 4**), suggesting a role for PNPLA3-I148M, either directly or indirectly, in restructuring this central hub of the secretory pathway and its interactome.

Dissecting the differences between WT PNPLA3 and PNPLA3-I148M can greatly enhance our understanding of the genetic basis for MASLD and fundamental lipid metabolism. As such, this work can help to refine our grasp of what “goes awry” with PNPLA3-I148M. The finding that PNPLA3 and PNPLA3-I148M are more enriched in the Golgi than the ER highlights the importance of the Golgi in lipid homeostasis and human disease. Many lipid metabolic enzymes localize to the ER (29, 48). The Golgi, though important for hepatic very-low-density lipoprotein (VLDL) biogenesis (49–51), has not been extensively investigated as a meaningful node for MASLD/MASH nor for PNPLA3 biology. As a bustling depot for intracellular cargo intimately integrated with the cytoskeleton, changes in the Golgi apparatus can have pleiotropic effects on signal transduction, lipid trafficking, and membrane homeostasis (52, 53). One study found that loss-of-function MASLD genetic risk factors introduced into HepG2 hepatoma cells, which express endogenous PNPLA3-I148M, caused enlarged Golgi cisternae relative to parental cells, but this study did not investigate the effects of PNPLA3-I148M alone on the Golgi (54). Lending credence to the findings presented here, an RNAi screen in *Drosophila* S2 cells identified members of the COPI vesicular trafficking machinery, which facilitate retrograde Golgi-to-ER and intra-Golgi vesicular transport, as significant regulatory factors for LD formation and morphology (55). Additionally, liver proteomics and phosphoproteomics in a diet-induced MASLD mouse model demonstrated reorganization of the secretory pathway, including redistribution of COPI proteins and segregation of Golgi proteins to LDs (56). Although we have established an association of PNPLA3-I148M with the Golgi, future work is needed to investigate the determinants of localization, considering that there are no known pathways for Golgi-to-LD protein trafficking.

Membrane contact sites have gained increasing attention as important nodes of inter-organelle communication and sites of ion, lipid, and metabolite transfer (47). A systems-level characterization of organelle contact sites in COS-7 cells identified LD interactions with most other organelle types, with Golgi contacts representing the second most abundant heterotypic LD interaction (∼ 15% of LDs contacted the Golgi) (57). Although LD-Golgi interactions are clearly significant and potentially important in fatty liver disease (56, 58), these contact sites are poorly characterized, in part due to the unique shape and composition of the Golgi and the diversity of Golgi cisternae within a single cell (59). Although we found closer apposition of LDs and the Golgi in I148M cells compared to WT cells, the nature of these interactions, including whether they are directly mediated by PNPLA3-I148M, is the topic of future work. Limitations in antibodies suitable for endogenous PNPLA3 immunofluorescence have made such detailed studies challenging to conduct thus far.

Another notable finding from this work is that PNPLA3 and PNPLA3-I148M can associate with membrane phosphoinositides despite lacking an annotated phosphoinositide-binding domain, such as the pleckstrin homology (PH) or FYVE domain (42). Phosphoinositides are important for regulating membrane dynamics and maintaining organellar identity, especially within the secretory and endolysosomal pathways (42). Phosphoinositides and phosphoinositide-binding proteins also facilitate organellar junctions and nonvesicular lipid trafficking between cellular compartments, exemplified by the role of oxysterol binding protein (OSBP) in coupling sterol trafficking from the ER to the Golgi with PtdIns(4)P transport from the Golgi to the ER, leading to its hydrolysis (60). An association between PNPLA proteins and phosphoinositides is not unprecedented, as host cell plasma membrane localization of the *Pseudomonas aeruginosa* virulence factor ExoU, a PNPLA family member, is mediated by its association with PtdIns(4,5)P2 (61). Future work will uncover the nature of the PNPLA3-phosphoinositide interaction and the role that these lipids play in PNPLA3 biogenesis and function.

The knock-in cell lines developed in this study can be used to delve deeper into PNPLA3 biology. Although there are limitations to using cell lines to study the role of proteins in human disease, the key differences between mouse and human PNPLA3 tissue distribution and primary sequence, coupled with technical limitations and heterogeneity of primary human hepatocytes, prompted us to develop a controlled hepatoma cell system to directly compare the WT and I148M proteins. The fact that PNPLA3-I148M drives the full spectrum of MASLD, including HCC, further supports use of this cellular system. One published study used genetically engineered human induced pluripotent stem cells to model PNPLA3-I148M function with similar reasoning to ours (15). However, those authors concluded that *PNPLA3-I148M* is a loss-of-function mutation rather than neomorphic, the former of which is inconsistent with both mouse and human data pointing to PNPLA3-I148M protein as a driver of MASLD. The characteristic disease-like changes we observed in Hep3B-PNPLA3-I148M cells bolsters our ability to extrapolate findings to the disease state. Accordingly, we confirmed our observation of altered LD-Golgi dynamics in primary human PNPLA3-I148M hepatocytes.

Introducing a single amino acid change into PNPLA3 can change the morphological, proteomic, and transcriptomic landscape of the cell. Lipid metabolic pathways are tightly integrated and regulated, and the downstream effects of PNPLA3-I148M on the cellular proteome are pleiotropic, much like MASLD. Unraveling the causes and effects of the phenotypes and identifying the most upstream function of PNPLA3-I148M, starting with its initial biogenesis, could greatly enhance our understanding of lipid homeostasis and lead to the development of impactful therapeutics to curb the silent epidemic that is MASLD.

## Materials and Methods

### Cell lines and plasmids

Hep3B cells (ATCC) were grown in Eagle’s Minimum Essential Medium (EMEM; ATCC) supplemented with 10% fetal bovine serum (FBS; Gibco) and 1x Antibiotic-Antimycotic (Gibco). Prior to gene editing, cells were grown in antibiotic-free medium. The endogenous *PNPLA3* locus in Hep3B cells was edited using CRISPR to produce a cell line that is homozygous for *PNPLA3-I148M*. First, ribonucleoprotein complexes (PNA Bio) were generated at room temperature for 10 min. ssDNA molecules were subsequently added to the mixture to generate the “master mix” for co-delivery into Hep3B cells (Table S1). Cells were washed in PBS and resuspended in Resuspension Buffer R (Invitrogen). Then, cells were mixed 1:1 (v/v) with the master mix for electroporation using a Neon Transfection System (Invitrogen; pulse voltage = 1600, pulse width = 20, pulse number = 1) and immediately transferred to fresh growth medium lacking antibiotics. After growth to a sufficient density, genomic DNA was isolated and editing was confirmed by restriction digestion and droplet digital PCR (Bio-Rad). Cell lines that were edited were clonally selected by limiting dilution. Clonally isolated PNPLA3-I148M cell lines were validated with Sanger sequencing, ddPCR, and next-generation sequencing.

To generate endogenously tagged cell lines, confirmed Hep3B-PNPLA3 (WT) and Hep3B-PNPLA3-I148M clones were edited to insert a 72-bp HA-HiBiT tag at the 3’-end of the endogenous *PNPLA3* locus. Gene editing was performed essentially as described above (Table S1). Following electroporation, cells were added to fresh medium lacking antibiotics but containing 20 μM of the NHEJ inhibitor NU7026 (Selleck Chemicals). Following 48-72 h incubation, cells were sorted (BD FACSMelody Cell Sorter) into 96-well plates for clonal selection. Sorting media (pre-filtered) contained 50% FBS, 40% conditioned media (from previous passage), and 10% fresh media supplemented with 10 μM Y-27632 (Sigma) and 1x insulin/transferrin/sodium selenite (ITS; Sigma). Following growth, colonies were selected and screened by TOPO PCR cloning and Sanger sequencing. Clonal identities were further confirmed by NGS-based amplicon sequencing.

pTT5-PNPLA3-His and pTT5-PNPLA3-I148M-His were generated by inserting the coding sequences into the pTT5 vector (62) using Golden Gate Assembly. Briefly, two gBlocks for each construct (Twist Bioscience) were designed with proper overhang sequences using Geneious software (Dotmatics). gBlocks and vector were digested and ligated (in a single reaction) with BsmBI (Thermo Scientific) and T4 DNA ligase (Thermo Scientific). TOP10 cells (Invitrogen) were transformed with the Golden Gate Assembly reactions and colonies were selected on LB-Agar plates containing carbenicillin (Teknova) for Sanger sequencing confirmation (Azenta Life Sciences) using primers pTT5_PNPLA3_fwd, pTT5_PNPLA3_rev, pTT5_PNPLA3_int1, and pTT5_PNPLA3_int2 (Table S2).

pTT5-PNPLA3-FLAG and pTT5-PNPLA3-I148M-FLAG were generated using the respective His-tagged construct vectors as templates. Q5 Site-Directed Mutagenesis (NEB) with primers PNPLA3-FLAG_fwd and PNPLA3-FLAG_rev (Table S2) was employed to insert the coding sequence for a FLAG tag and STOP codon before the sequence for the His tag.

For siRNA treatment, Silencer Select Negative Control #1 (Invitrogen 4390843) and Silencer Select PNPLA3 (Invitrogen s37253) siRNA reagents were used. Cells were reverse-transfected using Lipofectamine RNAiMAX (Invitrogen), following the manufacturer’s protocol, and treated for 72 h.

### Allele-specific TaqMan assay

To confirm PNPLA3 variant status (WT vs. I148M) of cell lines and primary hepatocytes, RNA was first isolated by the RNeasy Mini Kit (Qiagen). Samples were then analyzed by digital PCR using a QIAcuity Eight Platform System (Qiagen) with a custom-made allele-specific TaqMan probe ANTZ9CG (ThermoFisher Scientific) that specifically detects PNPLA3 message at the location of the nucleotide change in rs738409 and quantifies reference (WT) copies and mutant variant (I148M) copies. For reference, the pan-PNPLA3 TaqMan probe Hs00228747_m1 (ThermoFisher Scientific) was used. The housekeeping gene probe Hs00427620_m1 (TBP; ThermoFisher Scientific) was employed to normalize expression. Manufacturer’s protocols were followed for dPCR experiments.

### Immunoblotting

Unless otherwise specified, gel samples were prepared by boiling for 5 min in 2x SDS sample buffer (125 mM Tris, 4% SDS, 20% glycerol, 0.04% bromophenol blue, pH 6.78) supplemented with 5% β-mercaptoethanol (Sigma). Samples were briefly centrifuged prior to loading onto 4-12% NuPAGE Bis-Tris mini-gels (ThermoFisher Scientific). For HiBiT tag detection, proteins were transferred from the gel onto a 0.45 µm nitrocellulose membrane (ThermoFisher Scientific) at 100 V for 1 h in a Mini Trans-Blot Electrophoretic Transfer Cell (Bio-Rad) kept at 4°C. For non-HiBiT detection, proteins were transferred onto 0.45 µm polyvinylidene difluoride membranes (ThermoFisher Scientific) using the same conditions. For HiBiT detection, nitrocellulose membranes were washed for 2 x 30 min at room temperature in PBS-0.1% Tween-20 and then incubated with LgBiT protein overnight at 4°C, following the protocol of the Nano-Glo HiBiT Blotting System (Promega). For detection of other proteins, membranes were blocked for 1 h at room temperature in 5% non-fat milk/PBS-T. Membranes were subsequently incubated with primary antibodies overnight at 4°C (Table S3), followed by washes at room temperature (1 x 15 min, 2 x 5 min) and incubation with HRP-conjugated secondary antibodies (Bio-Rad) for 1 h. Following three washes at room temperature, membranes were treated with SuperSignal West Dura Extended Duration Substrate (ThermoFisher Scientific) and affixed to a development cassette. Blots were exposed to BioMax MR film (Carestream Health) in a dark room and developed in an SRX-101A film developer (Konica Minolta).

### *In vitro* translation

Full-length DNA constructs for the coding sequences of human PNPLA3-HA, PNPLA3-I148M-HA, PNPLA3Δ42-62-HA, mouse Ii-op (63), and mouse RAMP4-op (63) were ordered as gBlocks (IDT) with a 5’ T7 <u>promoter</u>-linker-<u>Kozak</u> site (5’-<u>TAATACGACTCACTATAGGG</u>AATATTCTTGTTC<u>CCACCATG</u>…). gBlocks were PCR-amplified with Q5 High-Fidelity DNA Polymerase (NEB) using the forward primer T7-Kozak-start-fwd and the reverse primers HA-PolyA-rev (for PNPLA3-HA constructs) or Op-PolyA-rev (for Ii-op and RAMP4-op; Table S4). Amplified constructs were used as inputs for reactions (final reaction concentration of 6.96 ng/µL) using the TnT Quick Coupled Transcription/Translation System (Promega). Reactions were performed as described for the TnT system. Briefly, the rabbit reticulocyte lysate master mix was supplemented with EasyTag L-^35^-Methionine (4% final reaction volume; PerkinElmer). Canine rough pancreatic microsomes (cRMs; final A_280_ ∼ 2.5-3.5), prepared as described (64), were supplemented co-translationally. Reactions were incubated on a benchtop Thermomixer (Eppendorf) at 30°C for 55 min. To stop the reactions, puromycin dihydrochloride (Gibco) was added (final concentration of 2.5 mM), and samples were placed on ice. For post-translational reactions, cRMs were added following puromycin supplementation, and reactions were incubated at 30°C for an additional 30 min. Samples (kept on ice) were added to an equal volume of 1x PSB (100 mM KOAc, 2 mM Mg(OAc)_2_, 50 mM HEPES, pH 7.4) with 250 mM sucrose. Samples were ultracentrifuged in thickwall ultracentrifuge tubes (Thermo Scientific) for 15 min at 200,000 x *g*, 4°C (Sorvall MTX 150 ultracentrifuge with an S120-AT3 rotor). Supernatants were carefully removed and placed on ice. Pellets were resuspended in 1x PSB with 250 mM sucrose, and then mixed well with 100 mM (final concentration) sodium carbonate, pH 11. Tubes were left on ice for 30 min followed by another round of ultracentrifugation. Supernatants were carefully removed and kept on ice (wash fraction) and pellets were resuspended in PKB (1% SDS, 0.1 M Tris, pH 8.0). 16% of each fraction (initial supernatant, wash, and pellet) was mixed with 2x SDS sample buffer supplemented with 5% β-mercaptoethanol. Samples were boiled for 5 min prior to loading onto NuPAGE 4-12% Bis-Tris gels (Invitrogen). After washing with water and then with 5% glycerol, gels were placed upside down on thick filter paper (Bio-Rad) and covered with saran wrap to dry in a SGD2000 slab gel dryer (ThermoFisher Scientific). Gels were exposed to film and developed as above.

cRM Proteinase K (Lonza) and endoglycosidase H (NEB) digestion were performed as previously described (65, 66). Preparation of trypsin-or mock-treated cRMs was also done as previously described (63).

### Preparation of ER-derived microsomes from cells

HEK239T cells (ATCC) were split into 10-cm dishes and grown overnight in DMEM supplemented with 10% FBS. The following day, cells were transfected with pSNAPf-EF1α-PNPLA3-HA-HiBiT (Azenta Life Sciences) using Lipofectamine 3000 and grown for an addition 24 h. Cells were washed once in cold PBS, detached in cold PBS and centrifuged to sediment. Cell pellets were stored at -80°C until ready for use. Prior to microsome preparation, pellets were thawed on ice, resuspended in 10 mM HEPES-KOH, pH 7.5, and incubated for 10 min on ice. The cells were then pelleted, resuspended in homogenization buffer (10 mM HEPES-KOH, pH 7.5, 10 mM KCl, 1.5 mM MgCl_2_, 5 mM EGTA, 250 mM sucrose) and passed through a 27 G syringe needle 20 times. The homogenate was subjected to serial centrifugations (600 x *g* for 10 min, 3000 x *g* for 10 min, 100,000 x *g* for 60 min) prior to resuspension in membrane buffer (10 mM HEPES-KOH, pH 7.5, 50 mM KOAc, 2 mM Mg(OAc)_2_, 1 mM DTT, 250 mM sucrose), as described (66). Proteinase K digestion was done the same way as for cRMs, described above.

### Label free imaging for lipid droplet dynamics

Hep3B cells were seeded into 96-well glass-bottom plates (MatTek Corporation) at a density of ∼ 30,000 cells per 80 µL of phenol red-free EMEM medium (Quality Biological) and incubated overnight at 37°C. Then, prior to imaging, media was exchanged for fresh phenol red-free media with or without oleic acid-BSA (Sigma). Plates were briefly centrifuged to prevent meniscus formation and placed on the pre-equilibrated (37°C, 5% CO_2_ with humidity) Nanolive CX-A 3D Cell Explorer microscope (Nanolive). After approximately 1 h, thermalization was complete, and time-lapse refractive index images were captured every 3 min for 24 h using 3×3 gridscan mode (a field view of 275 x 275 µm). Images were segmented and quantified using the Smart Lipid Droplet Assay ^LIVE^ Module in the Eve software (Version 1.9.2.1760; Nanolive).

### Cellular fractionations

Crude membranes and lipid droplets were isolated as described (67) or by commercial precipitation-based methods to enrich for organelle sub-fractions (Invent Biotechnologies).

### Protein production

Proteins were expressed in suspension HEK 293-EBNA1 cells, as described (62, 68). First, complexes of plasmid and PEI MAX (Polysciences) were generated by mixing the appropriate pTT5 expression vector with transfection reagent (ratio of 1 µg vector : 4 µg PEI MAX) in F17 Expression Medium (Gibco). Complexes were incubated for 15 min at room temperature. Plasmid-PEI solutions were added to 293-EBNA1 cells (2 x 10^6^ cells/mL) in Expression Medium [F17 Medium supplemented with 0.1% Kolliphor P 188 (Sigma), 6 mM L-glutamine (Gibco), 25 µg/mL G418 Sulfate (Gibco)]. Cells were shaken overnight at 36°C, 5% CO_2_, 85% humidity. The next day, an equal volume of Expression Medium was added to the culture, and cells were grown in the same incubation conditions for 72 h. Cells were harvested by centrifugation at 3,700 x *g* for 30 min.

For PNPLA3-His constructs, proteins were purified as described (21), except that whole cell lysate (instead of just the membrane fraction) was passed over Ni-NTA. For PNPLA3-FLAG constructs, proteins were purified from the membrane fraction, as described (20).

### Lipid binding assays

PIP Strips (Echelon) were blocked overnight in PBS-T with 3% fatty acid-free bovine serum albumin (GoldBio) at 4°C. The next day, blocking buffer was removed and replaced with blocking buffer supplemented with 2 – 6 µg/mL of purified proteins (PLC-δ1 PH was obtained from Echelon). Strips were incubated on a shaker at room temperature for 1 h, followed by 3 x 10 min washes in PBS-T. Strips were then incubated in blocking buffer containing secondary antibodies (Table S3) on a shaker at room temperature for 1 h. Following 3 x 10 min washes, strips were developed with SuperSignal West Dura Extended Duration Substrate and imaged on a ChemiDoc MP Imaging System (Bio-Rad).

### Sample preparation for electron microscopy

Cells were grown on 3 mm synthetic sapphire disks (Technotrade International) and treated with 100 µM oleic acid overnight. Cells were then pre-fixed with 3% glutaraldehyde, 1% paraformaldehyde, 5% sucrose in 0.1 M sodium cacodylate. The disks were rinsed with fresh cacodylate buffer containing 10% Ficoll, placed into brass planchettes (Ted Pella, Inc.), and rapidly frozen with an HPM-010 high-pressure freezing machine (Bal-Tec). The frozen samples were transferred under liquid nitrogen to cryotubes (Nunc) containing a frozen solution of 2.5% osmium tetroxide, 0.05% uranyl acetate in acetone. Tubes were loaded into an AFS-2 freeze-substitution machine (Leica Microsystems) and processed at -90°C for 72 h, warmed over 12 h to -20°C, held at that temperature for 6 h, then warmed to 4°C for 2 h. The fixative was removed, and the samples rinsed 4x with cold acetone, after which they were infiltrated with Epon-Araldite resin (Electron Microscopy Sciences) over 48 h. The sapphire disks with affixed cells were flat-embedded onto a Teflon-coated glass microscope slide and covered with a Thermanox coverslip (Electron Microscopy Sciences). Resin was polymerized at 60°C for 48 h and the sapphire disks were excised, leaving the cells as a monolayer within a resin wafer.

### Electron microscopy and Dual-Axis tomography

Cells were observed by light microscopy and appropriate regions were extracted with a microsurgical scalpel and glued to the tips of plastic sectioning stubs. Semi-thin (170 nm) serial sections were cut with a UC6 ultramicrotome (Leica Microsystems) using a diamond knife (Diatome). Sections were placed on formvar-coated copper-rhodium slot grids (Electron Microscopy Sciences) and stained with 3% uranyl acetate and lead citrate. Gold beads (10 nm) were placed on both surfaces of the grid to serve as fiducial markers for subsequent image alignment. Sections were placed in a dual-axis tomography holder (Model 2040, E.A. Fischione Instruments) and imaged with a Tecnai T12-G2 transmission electron microscope operating at 120 KeV (ThermoFisher Scientific) equipped with a 2k x 2k CCD camera (XP1000; Gatan, Inc.). Tomographic tilt-series and large-area montaged overviews were acquired automatically using the SerialEM software package (69, 70). For tomography, samples were tilted ± 62° and images collected at 1° intervals. The grid was then rotated 90° and a similar series taken about the orthogonal axis. Tomographic data was calculated, analyzed, and modeled using the IMOD software package (70, 71) on iMac Pro and Mac Studio M1 computers (Apple, Inc.).

### Immunofluorescence

Hep3B cells were grown in Nunc Lab-Tek Chambered Slides (ThermoFisher Scientific). Primary human hepatocytes were grown in Collagen I-coated chambered slides (ibidi). Cells were washed and fixed in 4% formaldehyde (Sigma), as previously described (72, 73). Fixed cells were washed twice with PBS and incubated in 50 mM ammonium chloride (Sigma) for 10 min at room temperature. After two PBS washes, samples were permeabilized in 0.1% Triton X-100 for 10 min at room temperature. Fixed and permeabilized cells were washed twice in PBS and blocked in 5% normal goal serum (Gibco)/1% BSA for 30 min at room temperature. Primary antibodies (Table S3) were added in 1% BSA, and chamber slides kept at 4°C overnight. Samples were washed in PBS (3 x 5 min at room temperature) and then incubated with secondary antibodies diluted in 1% BSA for 1 h at room temperature (in the dark). Following PBS washes (3 x 5 min at room temperature), samples were incubated with 2 µM Hoechst 33342 (ThermoFisher Scientific) and HCS LipidTOX Green Neutral Lipid Stain (ThermoFisher Scientific) for 30 min at room temperature. The solution was exchanged for PBS, and images were obtained on a SP8 confocal microscope (Leica) using a 63X glycerol immersion lens.

### Juxtaposition analysis

Raw confocal images were analyzed in Fiji (74), where LD and TGOLN2 signals were separately segmented using machine learning pixel classification from Labkit (75). Segmentation data were imported as surfaces into Imaris 10.0.0 (Oxford Instruments) and juxtaposition analysis was performed by measuring the distance between surfaces (surfaces within 30 nm of each other were considered juxtaposed).

### Mass spectrometry

Hep3B cells were grown in 6-well plates (500,000 cells/mL) and washed three times with PBS prior to harvesting. Cell pellets (1 x 10^6^ cells/pellet) were flash frozen in liquid nitrogen and stored at -80°C until ready for analysis. Pellets were thawed on ice and resuspended in 0.05% SDS/0.5 M TEAB. Following multiple rounds of pipetting and brief vortexing, samples were passed through a 23 G syringe needle 30 times on ice and sonicated. Lysates were clarified by centrifugation at 16,000 x g for 10 min at 4°C. Supernatants were transferred to new tubes and total protein was quantified using a Bradford Assay (Bio-Rad). 20 µg protein per sample was reduced with 3 mM TCEP (Sigma) and alkylated with 10 mM iodoacetamide (Sigma) in the dark. Samples were digested overnight at 37°C with LysC and trypsin. 5 µg peptides from each sample were labelled using the TMTpro reagents (ThermoFisher Scientific) for 2 hours, followed by quenching with 5% hydroxylamine for 15 minutes, pooling, and drying down. Labelled peptides were mixed and analyzed using two-dimensional liquid chromatography and tandem mass spectrometry, as previously described (76). The pooled samples were desalted followed by phosphopeptide enrichment through sequential use of the High-Select TiO2 and Fe-NTA phosphopeptide enrichment kits (Thermo Scientific). All native peptides not captured by the phosphopeptide enrichment kits were then separated by offline medium pH C4 peptide fractionation (Accucore 150-C4, 2.6um pore size, 150mm X 2.1mm, Thermo Scientific) using gradient mobile phase conditions as previously reported (76). Fractionated peptides were concatenated into 6 pooled fractions, then dried down and stored at −80 °C until mass spectrometry analysis.

Liquid chromatography-mass spectrometry (LC-MS) analysis was carried out on an EASY-nLC 1000 (ThermoFisher Scientific) coupled to an Orbitrap Eclipse Tribrid mass spectrometer (ThermoFisher Scientific). Concatenated native peptide fractions were resuspended in 15 µL 2% acetonitrile, 0.2% formic acid, and 3 – 4 µg peptides per concatenated sample were loaded onto a monolithic column (Capillary EX-Nano MonoCap C18 HighResolution 2000, 0.1 x 2000 mm, Merck) fitted with a silica coated PicoTip emitter (New Objective FS360-20-10-D) and separated over 300 min at a flow rate of 500 nL/min with the following gradient: 2–6% Solvent B (10 min), 6-40% B (260 min), 40-100% B (1 min), and 100% B (29 min). Solvent A consisted of 97.8% water, 2% acetonitrile, and 0.2% formic acid, and solvent B consisted of 19.8% water, 80% acetonitrile, and 0.2% formic acid.

MS1 spectra were acquired in the Orbitrap at 120K resolution with a scan range from 375-2000 m/z, an AGC target of 4e5, and a maximum injection rate of 50 ms in Profile mode. Features were filtered for monoisotopic peaks with a charge state of 2-7 and a minimum intensity of 2.5e4, with dynamic exclusion set to exclude features after 1 time for 60 seconds with a 5-ppm mass tolerance. HCD fragmentation was performed with collision energy of 32% after quadrupole isolation of features using an isolation window of 0.7 m/z, an AGC target of 5e4, and a maximum injection time of 86 ms. MS2 scans were then acquired in the Orbitrap at 50K resolution in Centroid mode with the first mass fixed at 110. Cycle time was set at 1 s.

Phosphopeptide fractions from the TiO2 and Fe-NTA enrichment procedures were each resuspended in 10 uL 2% acetonitrile, 0.2% formic acid, and 8 µL peptides per sample were loaded onto an Aurora 25cm x 75µm ID, 1.6µm C18 reversed phase column (Ion Opticks) and separated over 136 min at a flow rate of 350 nL/min with the following gradient: 2–6% Solvent B (7.5 min), 6-25% B (82.5 min), 25-40% B (30 min), 40-98% B (1 min), and 98% B (15 min). All MS/MS parameters were identical to those for the native peptide runs.

Proteomics data analysis was performed in Proteome Discoverer 2.4 (Thermo Scientific) using the Byonic search algorithm (Protein Metrics) and Uniprot human database. Byonic search parameters for native peptide fractions were as follows: fully Tryptic peptides with no more than 2 missed cleavages, precursor mass tolerance of 10 ppm and fragment mass tolerance of 20 ppm, and a maximum of 3 common modifications and 2 rare modifications. Cysteine carbamidomethylation and TMTpro addition to lysine and peptide N-termini were static modifications. Methionine oxidation and lysine acetylation were common dynamic modifications (up to 2 each). Methionine loss on protein N-termini, methionine loss + acetylation on protein N-termini, protein N-terminal acetylation, and phosphorylation of serine, threonine, and tyrosine were rare dynamic modifications (only 1 each). Percolator FDRs were set at 0.001 (strict) and 0.01 (relaxed). Spectrum file retention time calibration was used with TMTpro addition to peptide N-termini and lysines as static modifications. Reporter ion quantification used a co-isolation threshold of 50 and average reporter S/N threshold of 10. Normalization was performed on total peptide amount and scaling was performed on all average. Peptide and protein FDRs were set at 0.01 (strict) and 0.05 (relaxed), with peptide confidence at least high, lower confidence peptides excluded, and minimum peptide length set at 6.

Statistical analysis of protein abundances was performed as previously described (77). Protein-level output files from Proteome Discoverer 2.4 were used for analyses. A limma test, implemented in R, was used to determine significance between WT and I148M cell lines (n = 8 per group). *p*-values were corrected for multiple testing by a Benjamini-Hochberg procedure.

For analysis of differential enrichment of biological process keywords in I148M versus WT cell lines, proteins significantly (adj. *p*-value < 0.01) decreased or increased by >1.5 fold in the I148M vs. WT PNPLA3 cell lines were compared with those not significantly changing. Biological process keywords for these sets of proteins were obtained from UniProt and Fisher’s exact test was used to determine a *p*-value, which was Bonferroni corrected, for each keyword. Only biological process keywords annotated for at least 5 proteins in at least one of the sets of proteins were considered.

Similar analyses were performed for disease-specific terms. Protein associations for these terms were collected from the Comparative Toxicogenomics Database (CTD; (78)), DisGeNET (79), and (45).

A list of liver-specific genes was obtained from the Tissue-specific Gene Expression and Regulation (TiGER) database.

### RNA-seq

Hep3B cells were grown overnight in 6-well plates (seeded at 500,000 cells/mL). Enough plates were seeded to allow for 5 replicates of the following conditions: WT & I148M basal, WT & I148M 30 min, 1, 2, 4, and 24 h oleic acid (100 µM) treatment. Following the specified treatment, cells (1 x 10^6^ cells/well) were washed three times with PBS and lysed in Buffer RLT (Qiagen). Lysates were transferred to Eppendorf tubes on ice and stored at -80°C until ready for RNA purification. Library preparation from purified RNA, sequencing and analysis protocols have been described previously (80). To determine the changes of gene expression, differential expression analysis between WT and I148M samples at each time point was conducted using DESeq2 (1.26.1) (81). Gene expression fold-change and adjusted *p*-values were generated using default parameters.

## Acknowledgments

We thank Mikhal Hardy and Mike Ollmann for initial generation of genetically modified Hep3B cells; Spiros Garbis for mass spectrometry/proteomics guidance and support; Tom Nguyen, Jiamiao Lu, Chi-Ming Li, Oliver Homann, Xuan Liu, and Xin Luo for RNA-seq sample preparation and data analysis support; Bram Estes, Kevin Graham, and Katie Douglas for support with cloning and protein production; and Malia Potts and Simon Jackson for critical reads and discussions of manuscript drafts. The Proteome Exploration Laboratory was supported by NIH OD010788, NIH OD020013, the Betty and Gordon Moore Foundation through grant GBMF775, and the Beckman Institute at Caltech. The EM work was supported by NIH grant P50AI150464 and the Beckman Electron Microscopy Center at Caltech.

## Author Contributions

D.J.S., R.J.D., and I.C.R. designed research; D.J.S., L.L., B.L., and M.S.L. performed research; D.J.S., L.L., J.L.M., B.L., and M.S.L. analyzed data; J.X. and J.F. contributed new analytical tools; D.J.S. wrote the original manuscript; J.L.M., R.V., I.C.R., and R.J.D. reviewed and edited the manuscript; R.J.D., I.C.R., and R.V. advised on the research; R.J.D. and I.C.R. supervised the research.

## Footnotes

† As of June 2023, the American Association for the Study of Liver Diseases (AASLD) has renamed non-alcoholic fatty liver disease (NAFLD) as metabolic dysfunction-associated steatotic liver disease (MASLD) and non-alcoholic steatohepatitis (NASH) as metabolic dysfunction-associated steatohepatitis (MASH) (https://www.aasld.org/new-nafld-nomenclature).

