## Supporting Information for "The fatty liver disease-causing protein PNPLA3-I148M alters lipid droplet-Golgi dynamics"

**This PDF file includes:**

Figures S1 to S5
Tables S1 to S5
SI Reference

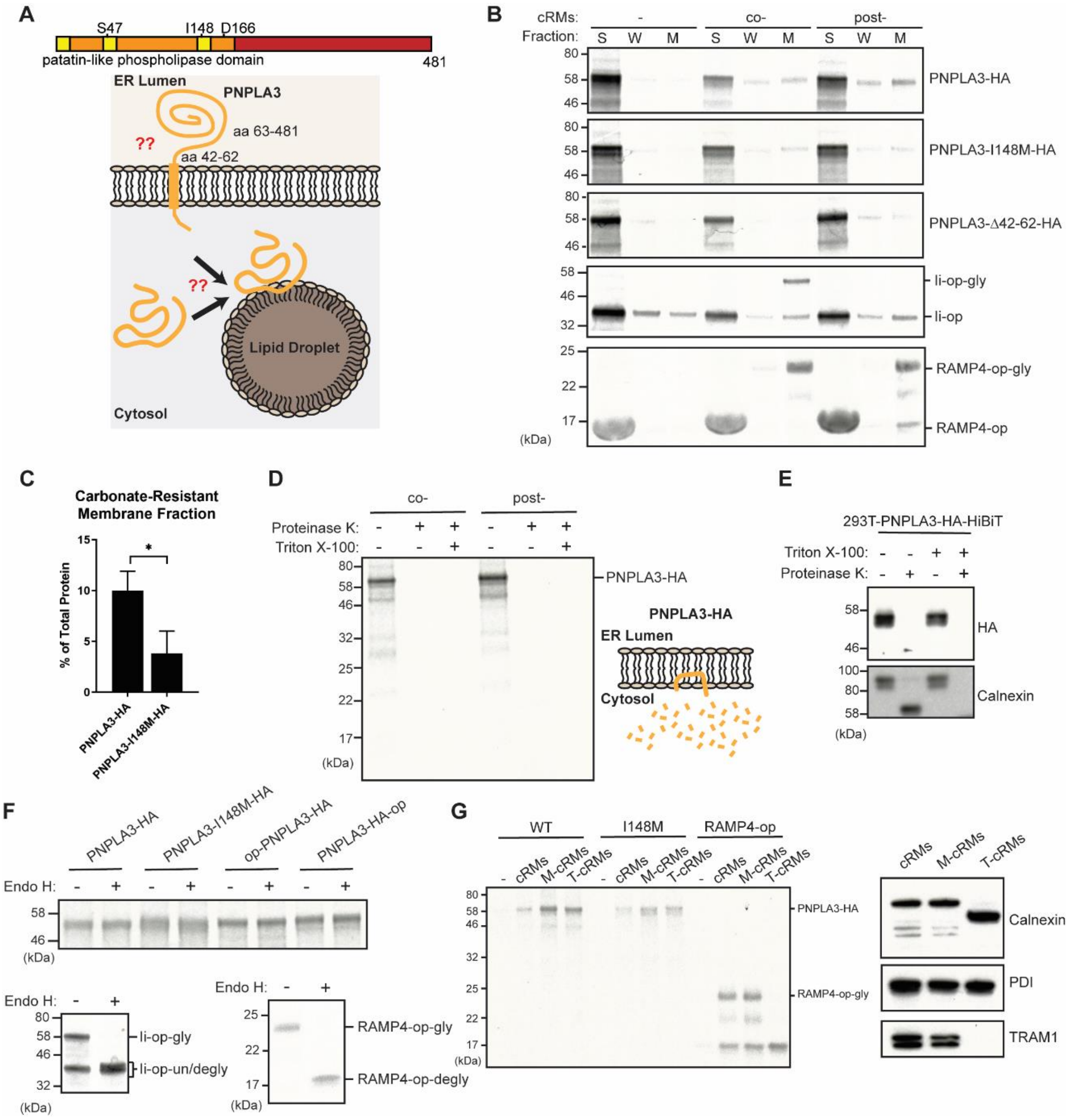


**Fig. S1. PNPLA3 is cytosolically exposed and lacks an ER-targeting signal sequence.** (A) Schematic (top) of the PNPLA3 primary sequence, with predicted hydrophobic segments of roughly ~ 20 amino acids in the PNPLA domain shown in yellow. Cartoon (bottom) depicts contrasting models of where PNPLA3 originates prior to localization on LDs, including a predicted ER topology. Amino acids 42-62 comprise a predicted type II signal anchor sequence. Residues S47 and D166 comprise the putative catalytic dyad. (B) PNPLA3-HA and variants, in addition to the control proteins invariant chain (Ii)-opsin tag and RAMP4-opsin tag, were translated in rabbit reticulocyte lysate in the presence of $^{35}$S-methionine. Canine rough microsomal membranes (cRMs) were added co- or post-translationally. Membranes were isolated, washed with sodium carbonate, then re-isolated. “S” = soluble fraction; “W” = carbonate wash; “M” = membrane fraction. (C) Quantification of PNPLA3 in membrane fractions. * $p < 0.05$, determined by Student’s t-test (n = 3). (D) cRMs added co- or post-translationally were incubated with Proteinase K with or without Triton X-100. (E) HEK293T cells were transiently transfected with a PNPLA3-HA-HiBiT vector. After 48 h, microsomes were isolated from the cells and subsequently treated with Proteinase K with or without Triton X-100. (F) cRM’s added post-translationally were solublized and treated with Endoglycosidase H. Control reactions were done with Ii-op and RAMP4-op. (G) cRM’s were mocktreated (M-cRMs) or trypsin-treated (T-cRMs) to remove exposed portions membrane proteins. These cRM's were added post-translationally to translation reactions. Following a carbonate wash and re-isolation of cRMs, localization of PNPLA3-HA constructs or RAMP4-op were assessed by SDS-PAGE (left). cRMs were characterized by immunonlotting to ensure trypsin digestion of exposed proteins (right).

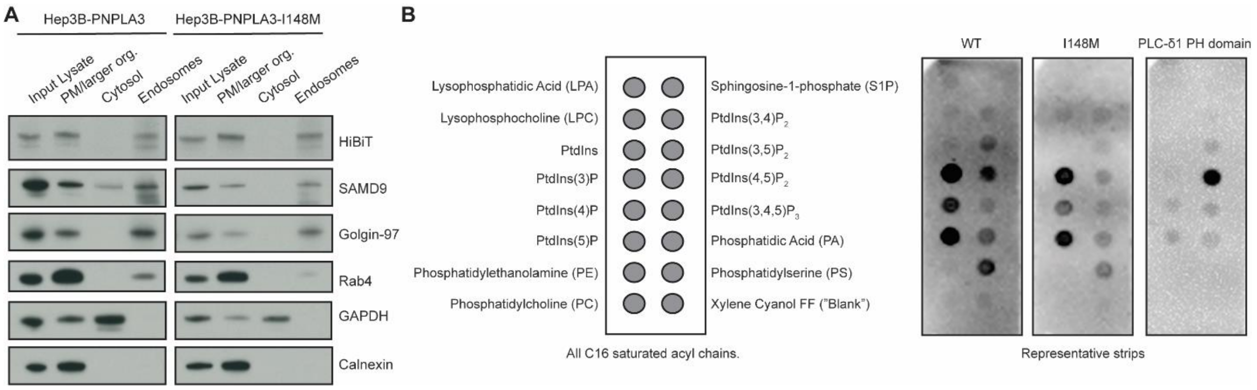


**Fig. S2. PNPLA3-I148M fractionates with endosomes and interacts with membrane phosphoinositides.** (A) Endogenous PNPLA3 and PNPLA3-I148M are enriched in the plasma membrane (PM)/larger organelle fraction and the endosomal fraction. (B) Purified PNPLA3-His, PNPLA3-I148M-His, and GST-PLC-δ1 PH domain were incubated with hydrophobic membranes spotted with 100 pmol of different lipids (left). After washing the membranes thoroughly, they were incubated with anti-His or anti-GST secondary antibodies (HRP-conjugated) and imaged using chemiluminescence. Experiment was repeated at least 3 times for each protein, with representative images shown (right).

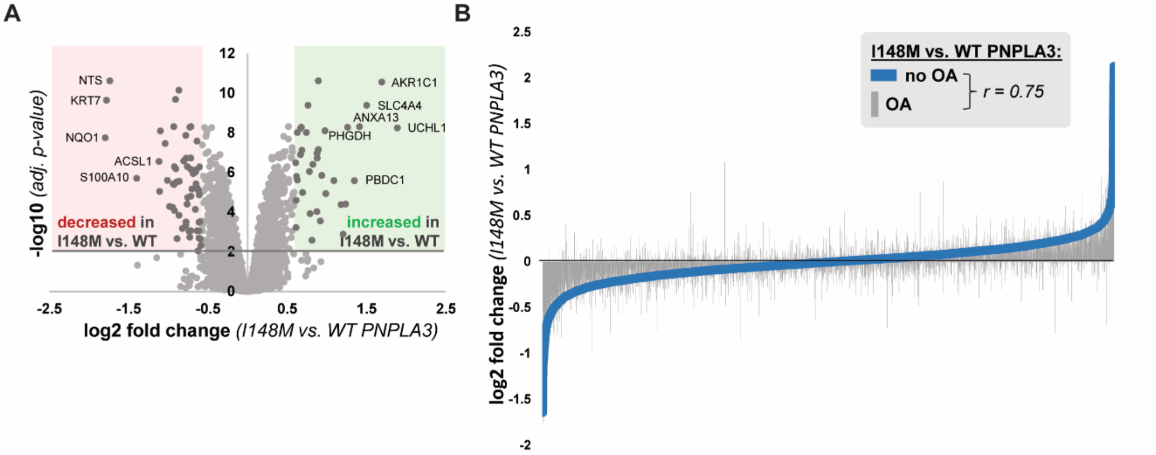


C

| Source | Disease | # Terms |
|---|---|---|
| Comparative Toxicogenomics Database (CTD) | fatty liver (MESH:D005234) | 1853 |
| Comparative Toxicogenomics Database (CTD) | hepatocellular carcinoma (MESH:D006528) | 1132 |
| DisGeNET | fatty liver disease (CUI: C4529962) | 741 |
| DisGeNET | liver fibrosis (CUI: C0239946) | 1179 |
| DisGeNET | adult hepatocellular carcinoma (CUI: C0279607) | 150 |
| Ref. 1 | liver-enriched genes in hepatocellular carcinoma | 188 |

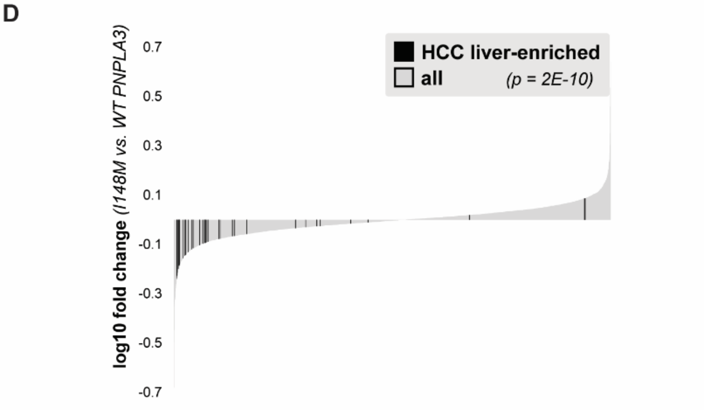


**Fig. S3. PNPLA3-I148M has a distinct effect on proteomic and transcriptomic cellular changes relative to oleic acid treatment.** (A) Volcano plot of proteins quantified by mass spectrometry in cells expressing WT or PNPLA3-I148M, following 16 h treatment with 200 µM oleic acid. Red and green compartments contain proteins significantly (adj. *p*-value < 0.01) decreased or increased, respectively, by >1.5 fold in the I148M vs. WT cell lines. (B) Comparison of the log2 fold change (I148M vs. WT PNPLA3) between two mass spectrometry experiments, in which cells were treated with or without oleic acid for 16 h. (C) Sources of liver disease-related gene sets. (D) Plot showing log10 fold change for proteins quantified in the I148M vs. WT PNPLA3 untreated cell lines. Those reported to be tissue-enriched genes in hepatocellular carcinoma (HCC) are shown in black (1). Kolmogorov-Smirnov *p*-value testing for a difference in fold change distribution between the two protein sets is reported.

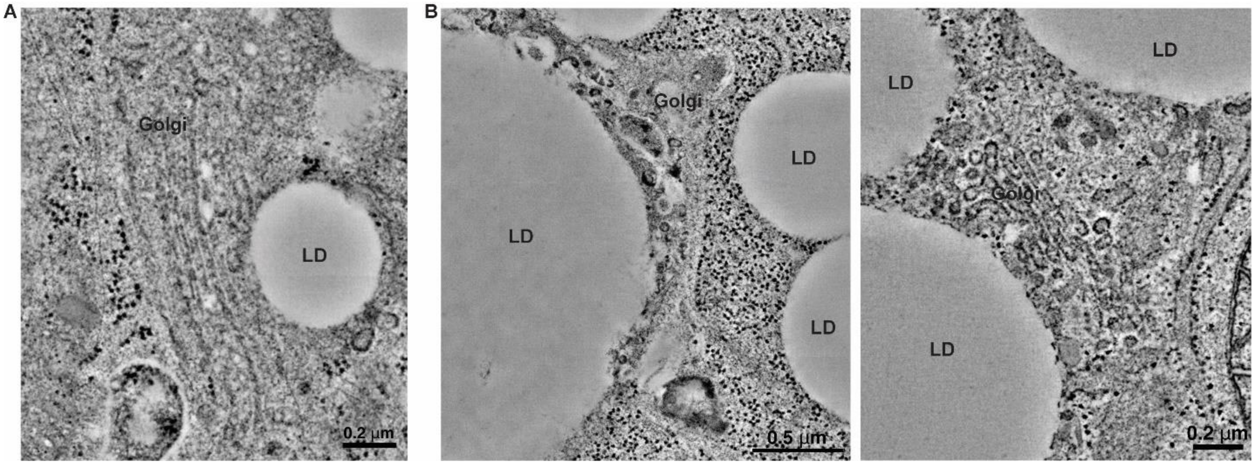


**Fig. S4. LD-Golgi contacts in Hep3B cells.** Representative TEM images from WT (A) and I148M (B) Hep3B cells pre-treated with 100 µM oleic acid for 16 h.

A

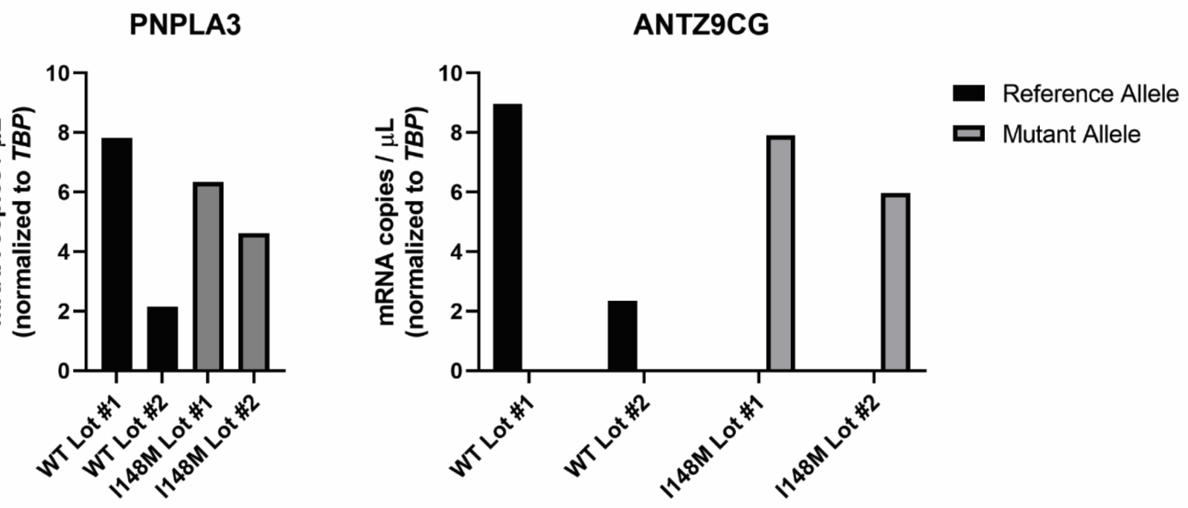


B

| Identifier | Vendor | Lot | BMI | Age at Death |
|---|---|---|---|---|
| WT #1 | Lonza | HUM4111C | 31.6 | 27 |
| WT #2 | Lonza | HUM4056B | 28.9 | 55 |
| I148M #1 | BioIVT | FZQ | 27.4 | 69 |
| I148M #2 | BioIVT | IXL | 25.0 | 56 |

**Fig. S5. Identification of primary human hepatocyte lots that express PNPLA3-I148M.** (A) Individual lots of primary human hepatocytes were assessed for *PNPLA3* expression by digital PCR using a TaqMan pan-PNPLA3 primer/probe set (left) and a custom TaqMan allele-specific primer/probe set (ANTZ9CG; right) that can distinguish the reference and mutant *PNPLA3* alleles. (B) Table showing key identifying information about primary human hepatocyte lots.

**Table S1. sgRNA and ssDNA for Hep3B gene editing.**

| Name | Description | 5'-3' Sequence |
|---|---|---|
| sgRNA-1 | For generation of *PNPLA3-I148M* | TGCTTCATCCCCTTCTACAG |
| sgRNA-2 | For generation of HA-HiBiT tag at 3' end | AGTCTGTGAGTCACTTGAGG |
| sgRNA-3 | For generation of HA-HiBiT tag at 3' end | AAGAGTCTGTGAGTCACTTG |
| ssDNA-1 | For generation of *PNPLA3-I148M* | CTTATGAAGGATCAGGAAAATTAAAAGGGTGCTCTCGCCTATAACTTCTCTCTCCTTTGCTTTCACAGGCCTTGGTATGTTCCTGCTTCATGCCCTTCTACAGTGGCCTTATCCCTCCTTCCTTCAGA |
| ssDNA-2 | For generation of *PNPLA3-I148M* | TAAAGTACCTGCTGGTGCTGAGGGGCTCTCCACCTTTCCCAGTTTTTCACTAGAGAAGAGTCTGTACCCATACGATGTTCCAGATTACGCTGGGAGCAGCGGGGTGAGCGGCTGGCGGCTGTTCAAGAAGATTAGCTGAGTCACTTGAGGAGGCGAGTCTAGCAGATTCTTTCAGAGGTGCTAAAGTTTCCCATCTTTGT |

**Table S2. Sequencing and Mutagenesis Primers.**

| Primer Name | 5'-3' Sequence | Description |
|---|---|---|
| pTT5_PNPLA3_fwd | CGAGGAGGATTTGATATTC | Sequencing pTT5-PNPLA3-His constructs |
| pTT5_PNPLA3_rev | CTTCCGAGTGAGAGACAC | Sequencing pTT5-PNPLA3-His constructs |
| pTT5_PNPLA3_int1 | AAGCAAGTTCCTCCGACAGG | Sequencing pTT5-PNPLA3-His constructs |
| pTT5_PNPLA3_int2 | AAGCTCAGTCTACGCCTCTG | Sequencing pTT5-PNPLA3-His constructs |
| PNPLA3_FLAG_fwd | CGACGATAAGTAGCACCATCA CCATCACCATCACCATTAG | Mutagenesis to make pTT5-PNPLA3-FLAG |
| PNPLA3_FLAG_rev | TCATCCTTGTAATCCCCGCTG CTCCCCAGACT | Mutagenesis to make pTT5-PNPLA3-FLAG |

**Table S3. Antibodies.**

| Antibody | Vendor | Catalog # | Use |
|---|---|---|---|
| β-actin | Cell Signaling | 4970 | Immunoblotting |
| GAPDH | EMD Millipore | MAB374 | Immunoblotting |
| α-tubulin | Cell Signaling | 2144 | Immunoblotting |
| PLIN2 | Cell Signaling | 95109 | Immunoblotting |
| Calnexin | Cell Signaling | 2679 | Immunoblotting |
| PERK | Cell Signaling | 3192 | Immunoblotting |
| TGN46 | Thermo Scientific | MA532523 | Immunoblotting |
| GM130 | Cell Signaling | 12480 | Immunoblotting |
| Golgin-97 | Cell Signaling | 13192 | Immunoblotting |
| SAMD9 | Sigma | HPA021319 | Immunoblotting |
| RAB4 | Cell Signaling | 2167 | Immunoblotting |
| COX IV | Cell Signaling | 4850 | Immunoblotting |
| GLUT1 | Cell Signaling | 12939 | Immunoblotting |
| TGOLN2 | Sigma | HPA012723 | Immunofluorescence |
| UCHL1 | Cell Signaling | 13179 | Immunoblotting |
| AKR1C1 | Thermo Scientific | PA521672 | Immunoblotting |
| PHGDH | Cell Signaling | 66350 | Immunoblotting |
| PDI | Enzo | ADISPA890D | Immunoblotting |
| TRAM1 | Thermo Scientific | PA521955 | Immunoblotting |
| HA-tag | Thermo Scientific | 26183 | Immunoblotting |
| Goat Anti-Rabbit IgG-HRP | Bio-Rad | 1706515 | Immunoblotting secondary Ab |
| Goat Anti-Mouse IgG-HRP | Bio-Rad | 1706516 | Immunoblotting secondary Ab |
| Anti-His-HRP | Thermo Scientific | MA121315HRP | Lipid binding assay |
| Anti-GST-HRP | Thermo Scientific | MA4004HRP | Lipid binding assay |
| Goat Anti-Rabbit IgG-AlexaFluor Plus 647 | Thermo Scientific | A32733 | Immunofluorescence secondary Ab |

**Table S4. Primers for *in vitro* translation.**

| **Primer Name** | **5’-3’ Sequence** |
|---|---|
| T7-Kozak-start-fwd | GAAATATAAGTAATACGACTCACTATAGGGAATATTCTTGTTCCCACCATG |
| HA-PolyA-rev | AATTTTTTTTTTTTTCAGGCGTAGTCGGGCACAT |
| Op-PolyA-rev | AATTTTTTTTTTTTTCAATCCACGGTTTTGTTGCTAAACGG |

**Table S5. Student's *t*-tests for lipid binding assays in Figure 3B.**

| | | *p*-value | |
|---|---|---|---|
| **Lipid 1** | **Lipid 2** | **WT** | **I148M** |
| PS | PtdIns(3)P | ns | ** |
| PS | PtdIns(4)P | * | *** |
| PS | PtdIns(5)P | ns | *** |
| PS | PtdIns(3,4)P2 | ns | * |
| PS | PtdIns(3,5)P2 | ns | * |
| PS | PtdIns(4,5)P2 | ns | *** |
| PtdIns(3)P | PtdIns | ns | * |
| PtdIns(3)P | LPA | ns | * |
| PtdIns(3)P | Sphingosine-1-phosphate | ns | * |
| PtdIns(3)P | No lipid | ns | ** |
| PtdIns(4)P | PtdIns(3,4)P2 | *** | ** |
| PtdIns(4)P | PtdIns(3,5)P2 | ** | ns |
| PtdIns(4)P | PtdIns(4,5)P2 | ns | * |
| PtdIns(4)P | PtdIns | *** | ** |
| PtdIns(4)P | LPA | *** | ** |
| PtdIns(4)P | LPC | *** | * |
| PtdIns(4)P | Sphingosine-1-phosphate | *** | ** |
| PtdIns(4)P | No lipid | *** | *** |
| PtdIns(5)P | PtdIns(3,4)P2 | * | ** |
| PtdIns(5)P | PtdIns(3,5)P2 | * | * |
| PtdIns(5)P | PtdIns(4,5)P2 | ns | ** |
| PtdIns(5)P | PtdIns | * | ** |
| PtdIns(5)P | LPA | * | ** |
| PtdIns(5)P | LPC | * | ** |
| PtdIns(5)P | Sphingosine-1-phosphate | * | ** |
| PtdIns(5)P | No lipid | * | *** |
| PtdIns(3,4)P2 | PtdIns(4,5)P2 | ns | * |
| PtdIns(3,4)P2 | No lipid | ns | * |
| PtdIns(3,5)P2 | No lipid | ns | * |
| PtdIns(4,5)P2 | PtdIns | ns | * |
| PtdIns(4,5)P2 | LPA | ns | * |
| PtdIns(4,5)P2 | Sphingosine-1-phosphate | ns | * |
| PtdIns(4,5)P2 | No lipid | ns | *** |

ns $p > 0.05$; * $p \leq 0.05$; ** $p \leq 0.01$; *** $p \leq 0.001$; **** $p \leq 0.0001$
